# Characterisation of the functional and transcriptomic effects of pro-inflammatory cytokines on human EndoC-βH5 beta cells

**DOI:** 10.1101/2022.11.29.518315

**Authors:** Caroline Frørup, Rebekka Gerwig, Cecilie Amalie Søndergaard Svane, Joana Mendes Lopes de Melo, Tina Fløyel, Flemming Pociot, Simranjeet Kaur, Joachim Størling

## Abstract

**Objective:** EndoC-βH5 is a newly established human beta-cell model which may be superior to previous models of native human beta cells. Exposure of beta cells to proinflammatory cytokines is a widely used in vitro model of immune-mediated beta-cell failure in type 1 diabetes and we therefore performed an in-depth characterisation of the effects of cytokines on EndoC-βH5 cells.

**Methods:** The sensitivity profile of EndoC-βH5 cells to the toxic effects of the pro-inflammatory cytokines interleukin-1β (IL-1β), interferon γ (IFNγ) and tumour necrosis factor-α (TNFα) was examined in titration and time-course experiments. Cell death was evaluated by caspase 3/7 activity, cytotoxicity, viability, TUNEL assay and immunoblotting. Mitochondrial function was evaluated by extracellular flux technology. Activation of signalling pathways and major histocompatibility complex (MHC) class I expression were examined by immunoblotting, immunofluorescence, and real-time quantitative PCR (qPCR). Glucose-stimulated insulin secretion (GSIS) and cytokine-induced chemokine secretion were measured by ELISA and Meso Scale Discovery multiplexing electrochemiluminescence, respectively. Global gene expression was characterised by stranded RNA sequencing.

**Results:** Cytokines increased caspase activity and cytotoxicity in EndoC-βH5 cells in a time- and dose-dependent manner. The proapoptotic effect of cytokines was primarily driven by IFNγ. Cytokine exposure caused impaired mitochondrial function, diminished GSIS, and induced secretion of chemokines. At the signalling level, cytokines increased the phosphorylation of signal transducer and activator of transcription 1 (STAT1) but not c-jun N-terminal kinase (JNK) and did not cause degradation of nuclear factor of kappa light polypeptide gene enhancer in B-cells inhibitor α (IκBα). MHC class I was induced by cytokines. Cytokine exposure caused significant changes to the EndoC-βH5 transcriptome including upregulation of *HLA* genes, endoplasmic reticulum stress markers, and non-coding RNAs. Among the differentially expressed genes were several type 1 diabetes risk genes.

**Conclusions:** Our study provides detailed insight into the functional and transcriptomic effects of cytokines on EndoC-βH5 cells. This knowledge will be helpful for future investigations studying cytokine effects in this cell model.

## Introduction

Cell models of human beta cells are important research tools for studying beta-cell biology and understanding beta-cell failure in diabetes. Most in vitro beta-cell research has been performed using immortalized rodent beta-cell lines such as rat INS-1 and mouse MIN6 [1–3]. Although the use of these models has advanced our understanding of several aspects of beta-cell biology under normal and pathophysiological conditions relevant for diabetes, the phenotypes of rodent beta-cells do not satisfactorily resemble native human beta cells. In 2011, Ravassard and colleagues successfully developed the first human beta-cell line, EndoC-βH1, derived from a human fetus [4]. For more than a decade, this cell line has proved a useful human beta-cell model although still with functional limitations compared to native human beta-cells including lower insulin production and secretory capacity, a tumorigenic phenotype and chronic infection with a xenotropic murine virus [5–9].

A new human beta-cell model, EndoC-βH5, has recently become commercially available (https://www.humancelldesign.com/). These cells demonstrate high insulin capacity and glucose responsiveness similar to native beta cells [10, 11]. However, several important aspects of the EndoC-βH5 cells still await to be characterised including their functional responses and behaviour under relevant diabetogenic conditions. For instance, it is highly relevant to establish the EndoC-βH5 cells’ sensitivity profile to pro-inflammatory cytokines which are mediators of beta-cell impairment in diabetes and constitute a commonly used in vitro model of type 1 diabetes pathogenesis [12–14]. In the present study, we therefore set out to thoroughly characterise the EndoC-βH5 cells’ response to cytokines at the functional, signalling, and transcriptomic levels.

## Methods

### Cell culture of EndoC-βH5

Frozen stocks of EndoC-βH5 cells were purchased from Human Cell Design (Toulouse, France). Upon thawing, cells had a viability between 74-91% (n=24). The EndoC-βH5 cells were seeded in appropriate plates coated with β-coat (Human Cell Design) and maintained in Ultiβ1 cell culture medium (Human Cell Design) containing 100 U/mL penicillin and 100 μg/mL streptomycin (Life Technologies). Based on the experimental setup, cells were seeded in 24-well (200,000 cells/well) or 96-well (50,000 cells/well) plates in duplicate and incubated in a humidified incubator at 37 °C with 5% CO2. Media change was carried out every 2-3 days. Cells were incubated for 0.5, 24, 48 and 96 hours in the presence or absence of recombinant human IL-1β (R&D systems), recombinant human IFNγ (PeproTech, Rocky Hill, NJ, USA) and/or recombinant human TNFα (R&D Systems).

### Cell death and viability

The ApoTox-Glo Triplex Assay (Promega, Madison, WI, USA) was used to measure live-cell protease activity, dead-cell protease activity and caspase-3/7 activity as markers of viability, cytotoxicity, and apoptosis, respectively. The assay was used according to the manufacturer’s protocol. Luminescence and fluorescence were measured on an Infinite M200 PRO plate reader (Tecan, Männedorf, Switzerland).

### Immunoblotting

Preparation of protein lysates, measurements of protein concentrations, and immunoblotting were performed as previously described [15]. Antibodies used were anti-cleaved caspase-7 (#9491S, Cell Signaling, 1:500), anti-MHC-I (#ALX-805-711-C100, Enzo, Farmingdale, NY, USA, 1:2000), anti-phosho-c-Jun N-terminal kinase (P-JNK) (#9252, Cell Signaling, 1:1000), anti-nuclear factor of kappa light polypeptide gene enhancer in B-cells inhibitor α (IκBα) (#J1512, Santa Cruz Biotechnology, Dallas, TX, USA, 1:500), anti-phospho-signal transducer and activator of transcription 1 (P-STAT1) (7649S, Cell Signaling, 1:1000), anti-Tubulin (T8203, Sigma-Aldrich, St. Louis, MO, USA, 1:2000), anti-GAPDH (#9482, Abcam, Cambridge, UK, 1:5000) and secondary HRP-conjugated anti-mouse (#7076) or anti-rabbit (#7074) (Cell Signaling) IgG antibodies. Tubulin or GAPDH was used as internal controls for normalisation.

### Immunofluorescence

Cells were seeded in black 96-well plates with clear bottom. After treatment for 48 hours, cells were fixed in 4% paraformaldehyde (Thermo Fisher) for 15 min at RT and permeabilized in 0.25% Triton X-100 (Sigma-Aldrich) for 20 min at RT. Blocking was done using 5% goat serum (Invitrogen) for 2 hours at RT. Staining of MHC-I (HLA-A/B/C molecules) was carried out using anti-MHC-I antibody (W6/32) (#ALX-805-711-C100, Enzo, 1:200 O/N at 4 °C) and Alexa Fluor 594-conjugated anti-mouse IgG antibody (#A11005, Invitrogen, 1:2000 for 2 hours RT). Nuclei staining was done in 0.5 μg/mL Hoechst 33324 (Sigma-Aldrich) in PBS. Furthermore, apoptosis was assessed by the detection of fragmented DNA using the Click-iT-Plus TUNEL Assay (Invitrogen, Waltham, Massachusetts, USA) according to the manufactures protocol. Fluorescence was measured using the ImageXpress PICO Automated Cell Imaging System (Molecular Devices (UK) Limited, Berkshire, UK) and analysed with the acquisition software (CellReporterXpress 2.9.3). Fluorescence was visualised using a Nikon ECLIPSE Ti2 microscope (Nikon, Tokyo, Japan). Images were acquired at x20 and x60 magnification and prepared using FIJI software (version 1.49).

### Chemokine measurements

The chemokines IFN-gamma-inducible protein 10 (IP-10)/C–X–C motif chemokine 10 (CXCL10), Thymus and activation-regulated chemokine (TARC)/CCL17, Eotaxin/CCL11, Eotaxin-3/CCL26, Macrophage Inflammatory Proteins (MIP)-1β/CCL3, MIP-1α/CCL4, IL-8/CXCL8, Monocyte chemoattractant protein (MCP)-1/CCL2, MCP-4/CCL13, and Macrophage-derived chemokine (MDC)/CCL22) were measured in the cell culture medium by the V-PLEX Chemokine Panel 1 Human Kit (Meso Scale Diagnostics LLC., Rockville, Maryland) using a MESO Quickplex SQ 120 instrument (Meso Scale Diagnostics).

### Glucose-stimulated insulin secretion

Cells were maintained in culture medium with or without cytokines for 48 hours prior to glucose-stimulated insulin secretion (GSIS), including 24 hours starvation in Ulti-ST starvation medium (Human Cell Design). Cells were washed twice and incubated for 60 min in βKrebs (Human Cell Design). Cells were incubated for 40 min in βKrebs supplemented with 0 or 20 mM glucose. Supernatants were collected and analysed for insulin. The cells were lysed in Cell Death Detection Lysis Buffer (Roche, Basel, Switzerland) for analysis of insulin content. Insulin was measured by ELISA (Mercodia, Uppsala, Sweden) and read on an Infinite M200 PRO plate reader (Tecan). Insulin was normalised to DNA content using CyQuant (Thermo Fisher Scientific) according to the manufacturer’s protocol.

### Seahorse extracellular flux

Cells were seeded in pre-coated XFe96 Seahorse plates (40,000 cells/well). After cytokine exposure, cells were washed twice and transferred into 180 μl Krebs-Ringer-HEPES (KRH) buffer (135 mM NaCl, 3.6 mM KCl, 10 mM HEPES, 0.5 mM MgCl2, 1.5 mM CaCl2, 0.5 mM NaH2PO4, 2 mM glutamine, 0,1% (w/v) fatty acid free BSA (Sigma-Aldrich), pH adjusted to 7.4 at 37 °C with NaOH). Cells were incubated for 1 hour at 37 °C in a CO2-free incubator prior to the experiment. After basal measurements, 11 mM glucose was injected to record respiratory response. Thereafter, 2 μM oligomycin (Sigma-Aldrich) was injected to inhibit ATP synthase and determine respiratory efficiency. Finally, 2 μM rotenone (Sigma-Aldrich), 2 μM antimycin A (Sigma-Aldrich) and 2.5 μM Hoechst 33324 was injected to inhibit mitochondrial respiration and stain for cell nuclei. Cell plates were immediately scanned using an ImageXpress PICO for cell count and quantified using the CellReporterXpress software. All data was normalised to cell count and corrected for non-mitochondrial respiration.

### RNA sequencing

RNA-seq libraries were sequenced on a DNBseq platform to produce 100 bp long paired-end reads. An average of 45-50 million reads per replicate (EndoC-βH5 n=4) were obtained after pre-processing and QC (removal of low-quality reads and adapter sequences) using SOAPnuke software [16]. Reads were aligned using TopHat (version 2.1.1) [17] to GChr38 genome with default parameters. Afterwards, reads were assigned to Ensembl version 107 gene annotation using htseq-count (version 0.9.1) [18] with default parameters. Minimum gene expression was defined based on a CPM cut-off of 0.5 in at least 4 samples. Differential gene expression analysis was performed on the two groups using the QL F-test in edgeR. Differentially expressed mRNAs and long non-coding RNAs (lncRNAs) were identified using a cut-off of abs(log2FC) >0.5 and an FDR-adjusted p-value <0.05. We extracted the gene expression of pinpointed candidate genes from a total of 152 loci recently shown to be associated with type 1 diabetes [19]. All statistical analyses were conducted using Bioconductor packages in R.

### Pathway analysis

Core analysis was performed using the Ingenuity Pathway Analysis (IPA) software (Qiagen, Valencia, CA, USA) on the differentially expressed genes to predict mechanistic and functional gene relationships, and identify top canonical pathways and molecular and cellular functions [20]. The summarized upstream regulators, diseases, and mechanisms for the differentially expressed genes were visualized as a network and grouped using the IPA Path Designer Module.

### Real-time qPCR

Gene expression levels of mRNAs and lncRNAs of interest as regulators of beta-cell death and cellular impairment [21, 22] were investigated by real-time quantitative PCR (qPCR). RNA was extracted using the RNeasy Mini Kit (Qiagen) and cDNA was synthesised using the iScript cDNA Synthesis Kit (Bio-Rad, Hercules, CA, USA). Real-time qPCR was carried out with TaqMan Assays and TaqMan Gene Expression Master Mix (Applied Biosystems, Waltham, MA, USA) using a CFX384 C1000 Thermal cycler (Bio-Rad). Relative expression levels were normalised to *GAPDH* as internal control [23].

### Statistical analysis

Data are presented as means ± SEM. Graphs were constructed using GraphPad Prism 9. Statistical analyses were performed using one- or two-way ANOVA and/or one- or two-tailed paired t-test where appropriate. P-values <0.05 were considered statistically significant. Multiple testing was corrected using the Benjamin-Hochberg’s correction.

## Results

### Cytokine sensitivity profile of EndoC-βH5 cells

To determine the sensitivity profile of the EndoC-βH5 cells, we initially examined the effects of cytokine exposure for 24, 48 and 96 hours, using cytokine concentrations previously used by us and others on isolated human islets and EndoC-βH1 cells (50 U/mL of IL-1β, 1000 U/mL IFNγ and 1000 U/mL TNFα) [24, 25]. Caspase-3/7 activity and cytotoxicity were significantly increased ~2-fold after 96 hours of cytokine exposure (Fig. 1a-b). The viability of the cells was slightly reduced, although not significantly (Fig. 1c).

**Figure 1:**
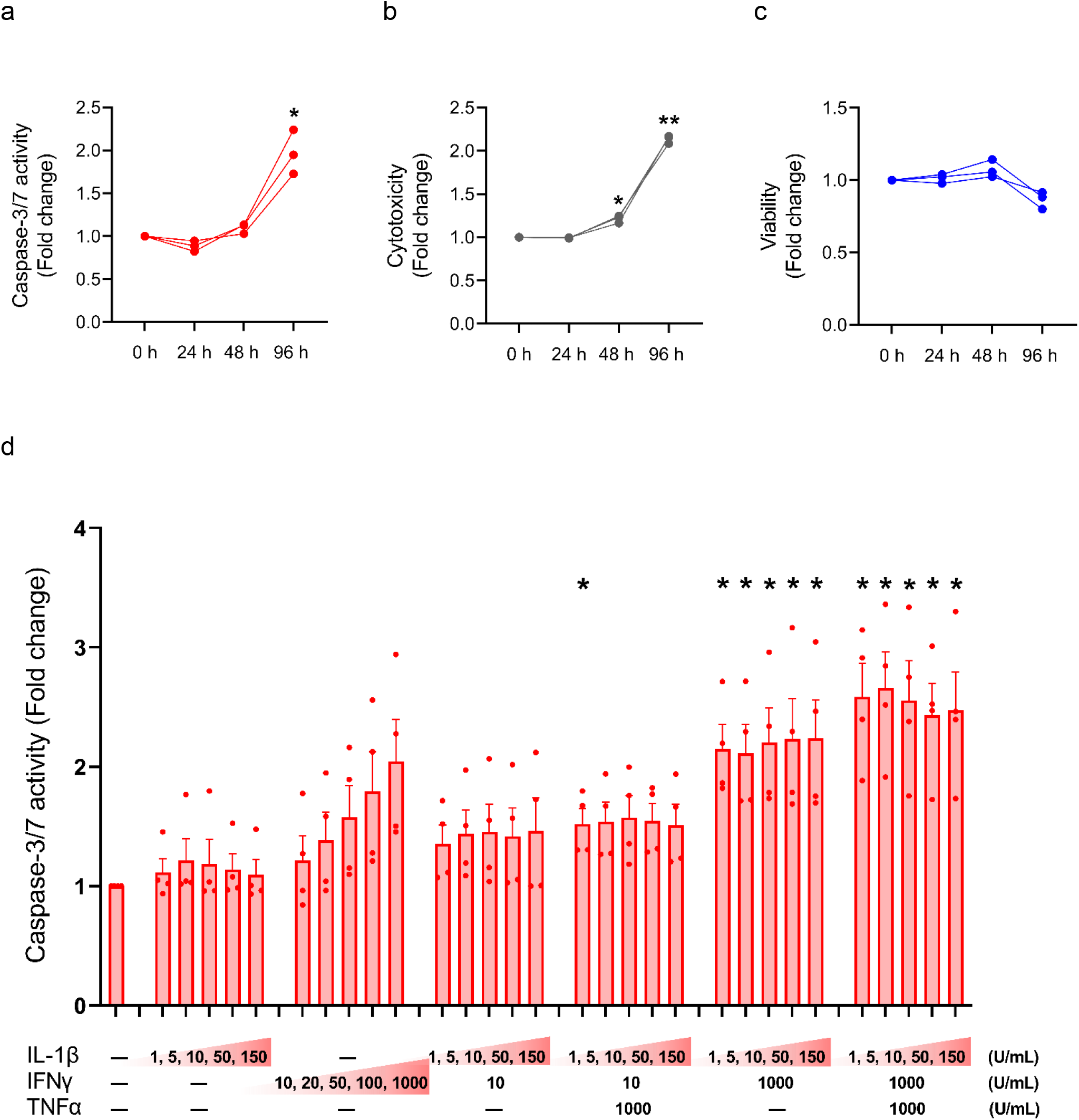
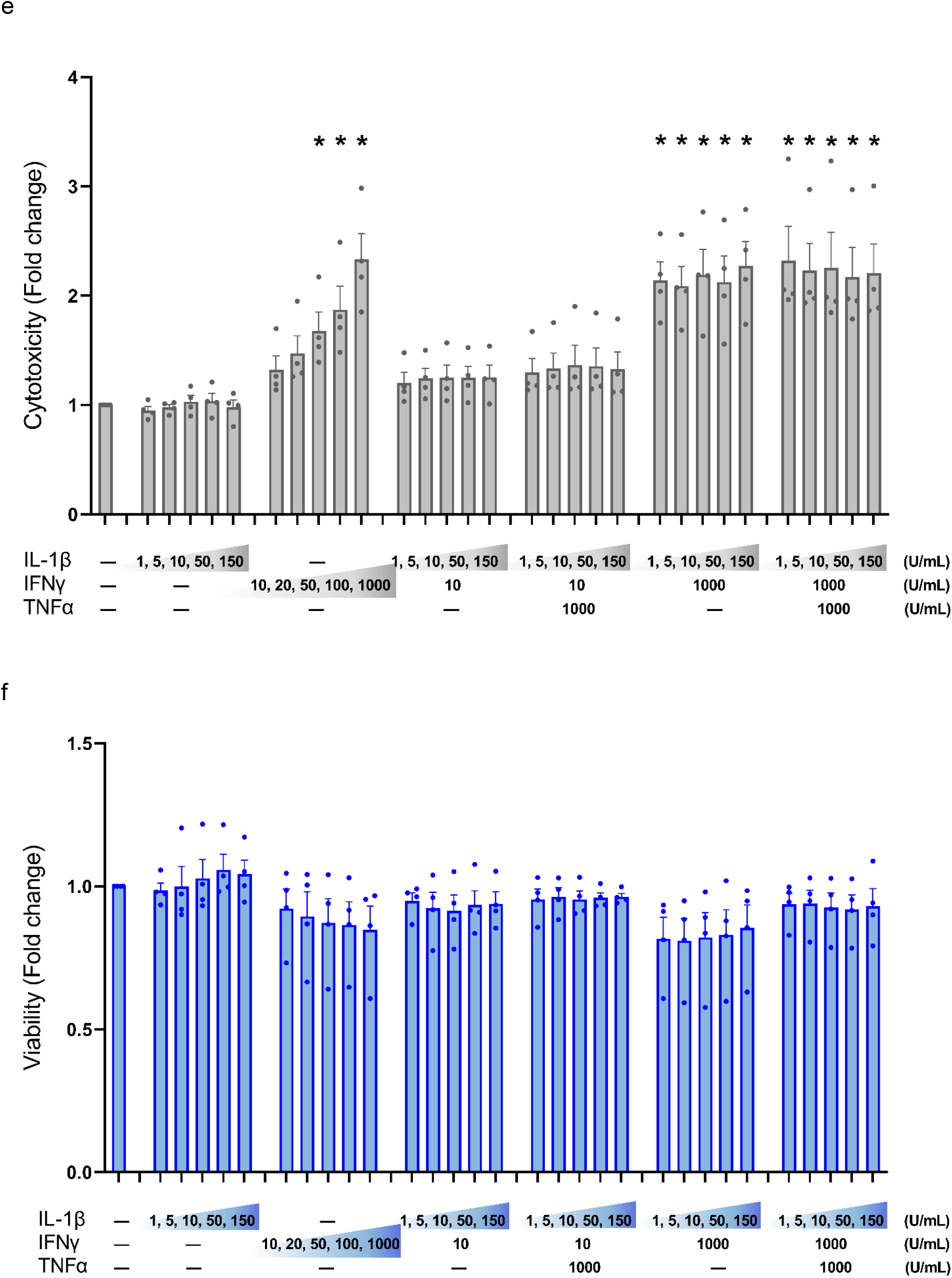
Cytokine sensitivity profile of EndoC-βH5 cells. (a) Caspase 3/7 activity, (b) cytotoxicity and (c) viability after exposure to cytokines (50 U/mL of IL-1β, 1000 U/mL IFNγ and 1000 U/mL TNFα) for 0 h, 24 h, 48 h and 96 h. Data are means ± SEM (n=3), *p<0.05; **p<0.01. (d) Caspase-3/7 activity, (e) cytotoxicity, and (f) viability after exposure to IL-1β, IFNγ and TNFα at the indicated concentrations for 96 h. Statistical significance is shown vertically above each bar. Data are means ± SEM (n=4). Adjusted p-values *p<0.05.

Next, we studied the responses to various doses of individual and combinations of cytokines after 96 hours of exposure. As seen in Fig. 1d and e, caspase-3/7 activity and cytotoxicity were dose-dependently induced by IFNγ alone. In contrast, IL-1β had no effects. When combined with IFNγ, neither IL-1β nor TNFα had synergistic effects on IFNγ-induced cell death. The reduction in cell viability by cytokine exposure did not reach statistical significance (Fig. 1f). These results indicate that only IFNγ exerts death-promoting and toxic effects on EndoC-βH5 cells.

End-stage apoptosis was evaluated by DNA fragmentation labelling with TUNEL. After 96 hours of cytokine exposure, end-stage apoptosis was elevated by ~1.6-fold in cells treated with either IFNγ alone or a combination of IL-1β, IFNγ and TNFα (Fig. 2). We observed around 20% TUNEL-positive cells in the untreated control condition indicative of a relatively high basal apoptosis rate (Fig. 2).

**Figure 2:**
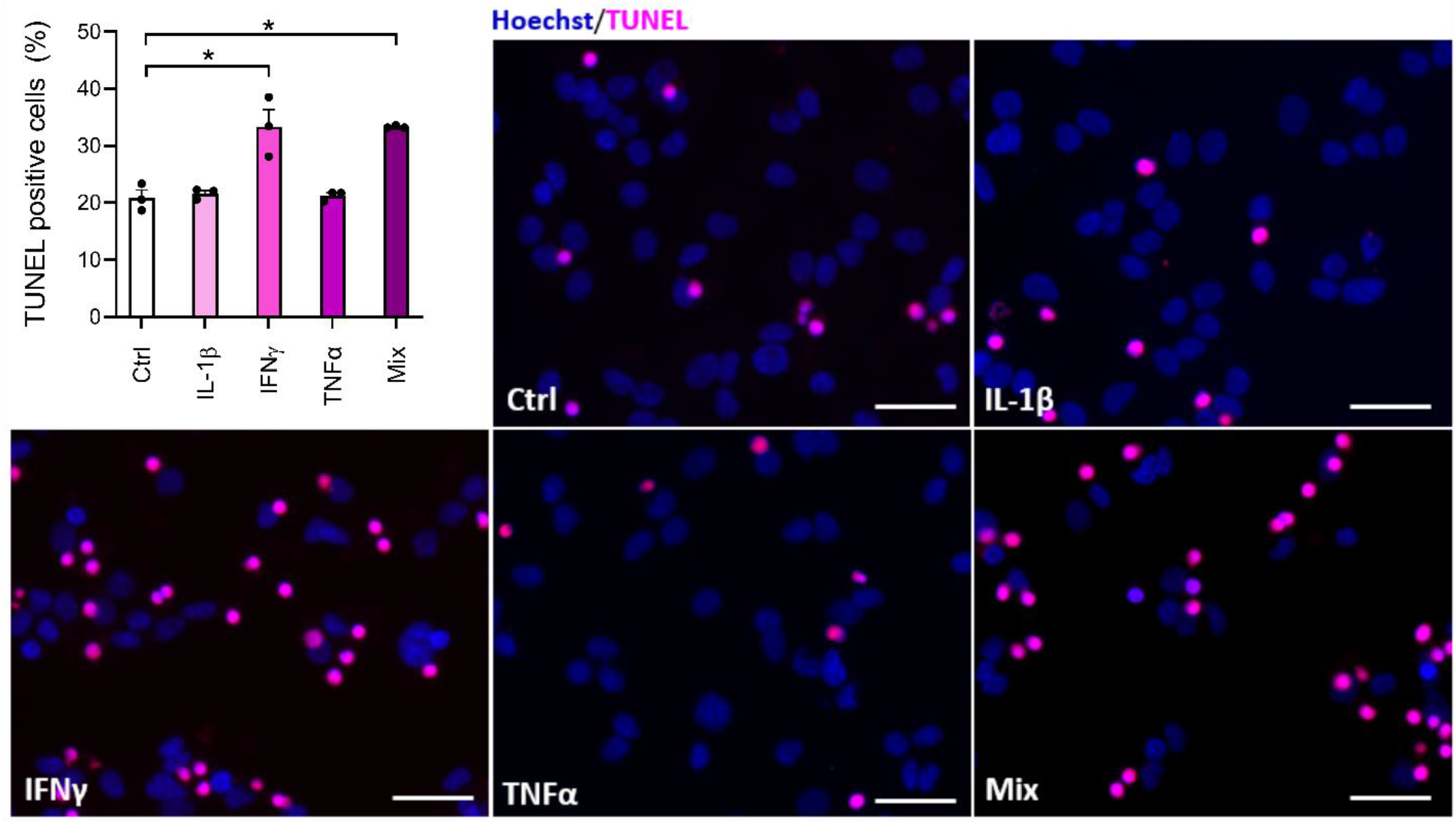
Cytokine-induced apoptosis in EndoC-βH5 cells. TUNEL assay of EndoC-βH5 cells after exposure to cytokines (50 U/mL IL-1β, 1000 U/mL IFNγ and 1000 U/mL TNFα) alone or in combination (Mix) for 96 h. Representative immunofluorescent images of TUNEL (pink) and nuclei staining with Hoechst 33324 (blue). Scalebar 50 μm (x20). Data are presented as % TUNEL positive to total nuclei count, with means ± SEM (n=3). *p<0.05.

### Cytokine signal transduction in EndoC-βH5 cells

To study cytokine signal transduction in EndoC-βH5 cells, we investigated key signalling pathways of each cytokine. We first confirmed that cytokine exposure leads to apoptotic caspase activation shown by immunoblotting of cleaved, activated caspase 7 (Fig. 3a). We then looked at proximal cytokine signal transduction by analysing P-STAT1 as a marker of IFNγ signalling, and P-JNK and IκBα as markers of IL-1β and TNFα signalling. Cytokine treatment for 30 min strongly enhanced P-STAT1, whereas no activating effects were seen on P-JNK and IκBα (Fig. 3b). INS-1E cell lysates were used as positive control and showed degradation of IκBα and elevated P-STAT1 and P-JNK after 30 min of cytokine exposure (Fig. 3b). A distal signalling event in cytokine-exposed beta-cells is the induction of endoplasmic reticulum (ER) stress. Individual cytokine exposure showed that only IFNγ induced ER stress as revealed by increased expression of the ER stress markers *DDIT3/CHOP* and *HRK/DP5* (Fig. 3c-d). Furthermore, IFNγ induced expression of *CXCL10* (Fig. 3e). These results suggest that only IFNγ is capable of inducing intracellular signalling in EndoC-βH5 cells.

**Figure 3:**
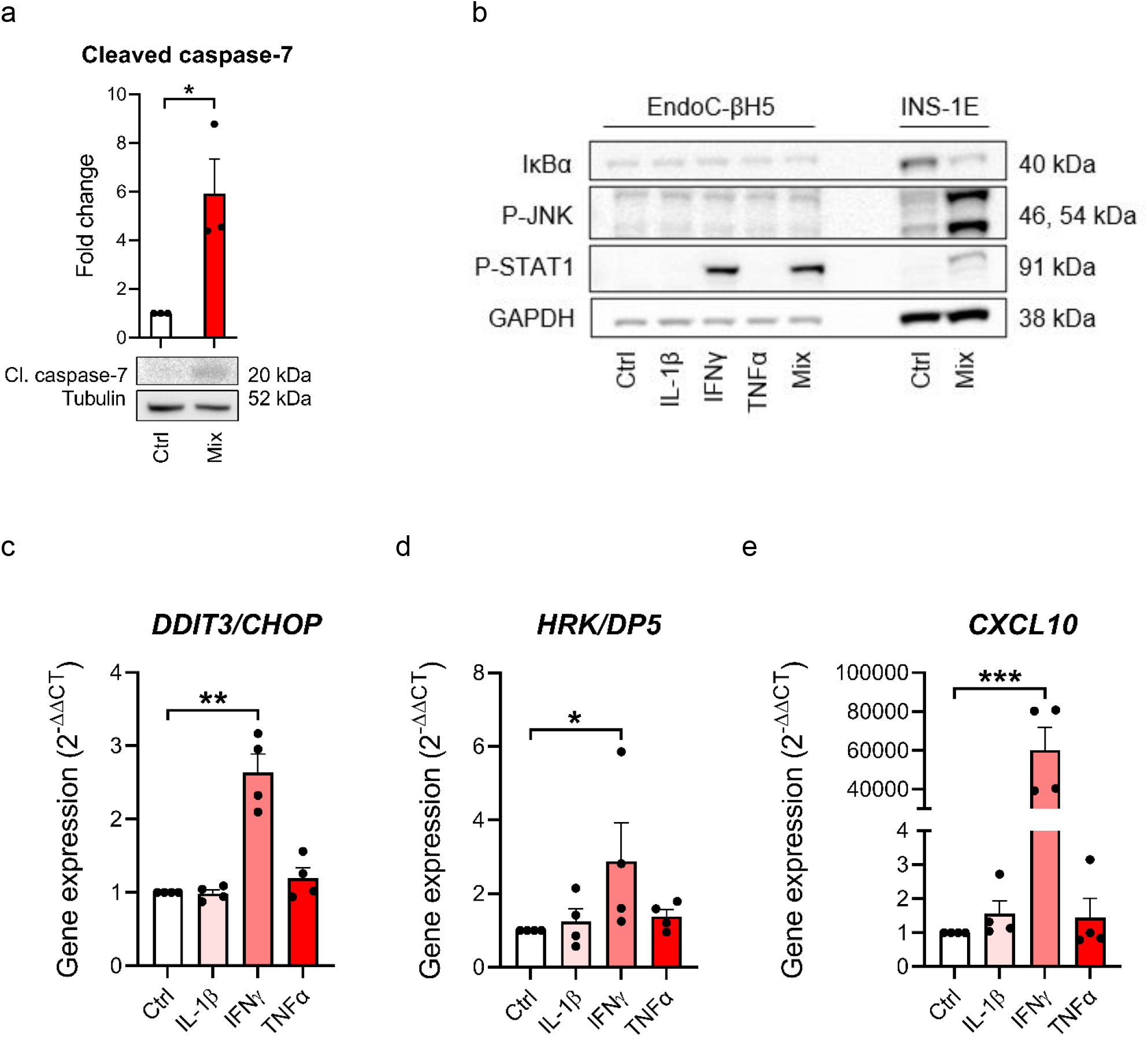
Cytokine-induced signal transduction in EndoC-βH5 cells. (a) Immunoblotting of cleaved caspase 7 following exposure to cytokines (50 U/mL IL-1β, 1000 U/mL IFNα and 1000 U/mL TNFα) (Mix) for 96 h. Tubulin was used as loading control (n=3). (b) Immunoblotting of IκBα, phosphorylated STAT1, and phosphorylated JNK following exposure to cytokines (50 U/mL IL-1β, 1000 U/mL IFNγ and 1000 U/mL TNFα) alone or in combination (Mix) for 30 min. GAPDH was used as loading control. Representative image (n=3). Lysates from INS-1E cells with/without cytokine exposure were used as positive control. Gene expression of (c) *DDIT3/CHOP,* (d) *HRK/DP5* and (e) *CXCL10* in cells exposed to individual cytokines for 48 h. *GAPDH* was used as housekeeping gene. Data are means ± SEM (n=4), *p<0.05, **p<0.01, ***p<0.001.

### MHC-I expression in EndoC-βH5 cells

We next investigated MHC class I (MHC-I), a hallmark of immune-mediated beta-cell destruction in type 1 diabetes, in the EndoC-βH5 cells in response to cytokines. By immunoblotting, protein expression of MHC-I was elevated after 96 hours of cytokine treatment (Fig. 4a). Consistent with this, cytokines upregulated the expression of *HLA-A/B/C* which encode MHC-I (Fig. 4b-d). These effects were driven primarily by IFNγ alone (Fig. e-f). Induction of MHC-I was confirmed by immunofluorescence (Fig. 4h). These findings demonstrate that cytokines upregulate MHC-I in EndoC-βH5 cells.

**Figure 4:**
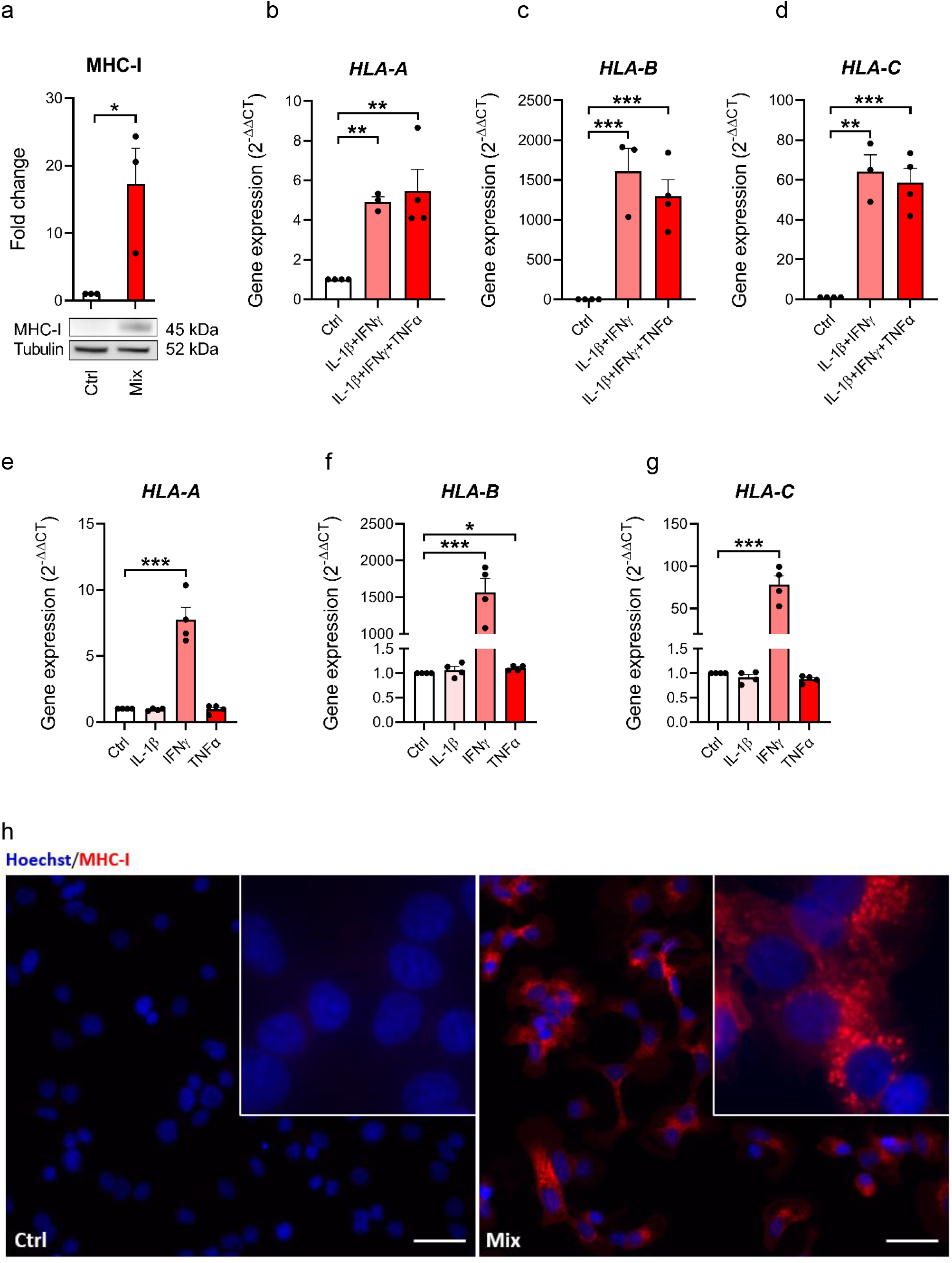
Cytokine-induced upregulation of MHC-I in EndoC-βH5 cells. (a) Immunoblotting of MHC-I in cells exposed to cytokines (50 U/mL IL-1β, 1000 U/mL IFNα and 1000 U/mL TNFα) (Mix) for 96 h. Tubulin was used as a loading control. Data are means ± SEM (n=3). Gene expression of *HLA-A* (b) *HLA-B* (c) and *HLA-C* (d) in cells exposed to 2 or 3 cytokines for 48 h, and gene expression of *HLA-A* (e) *HLA-B* (f) and *HLA-C* (g) in cells exposed to individual cytokines for 48 h. *GAPDH* was used as housekeeping gene. Data are means ± SEM (n=3-4). (h) Immunofluorescence staining of MHC-I (red) and Hoechst 33324 nuclei staining (blue) in EndoC-βH5 cells after 48 h of cytokine exposure. Scalebar 50 μm (20x) and zoom (60x). Representative images are shown. *p<0.05, **p<0.01, ***p<0.001.

### Secretion of chemokines from EndoC-βH5 cells

We examined if cytokines increase secretion of chemokines which play important roles in the immune cell – beta cell crosstalk in type 1 diabetes. The accumulated levels of 10 chemokines were measured in the cell culture medium after 48 hours of exposure to IL-1β, IFNγ and TNFα in combination or alone. All chemokines measured were increased in response to the treatment with the cytokine mixture (Fig. 5a). For treatment with the individual cytokines, IFNγ was the sole cytokine capable of increasing chemokine secretion (Fig. 5b). These data show that EndoC-βH5 cells secrete chemokines upon treatment with cytokines.

**Figure 5:**
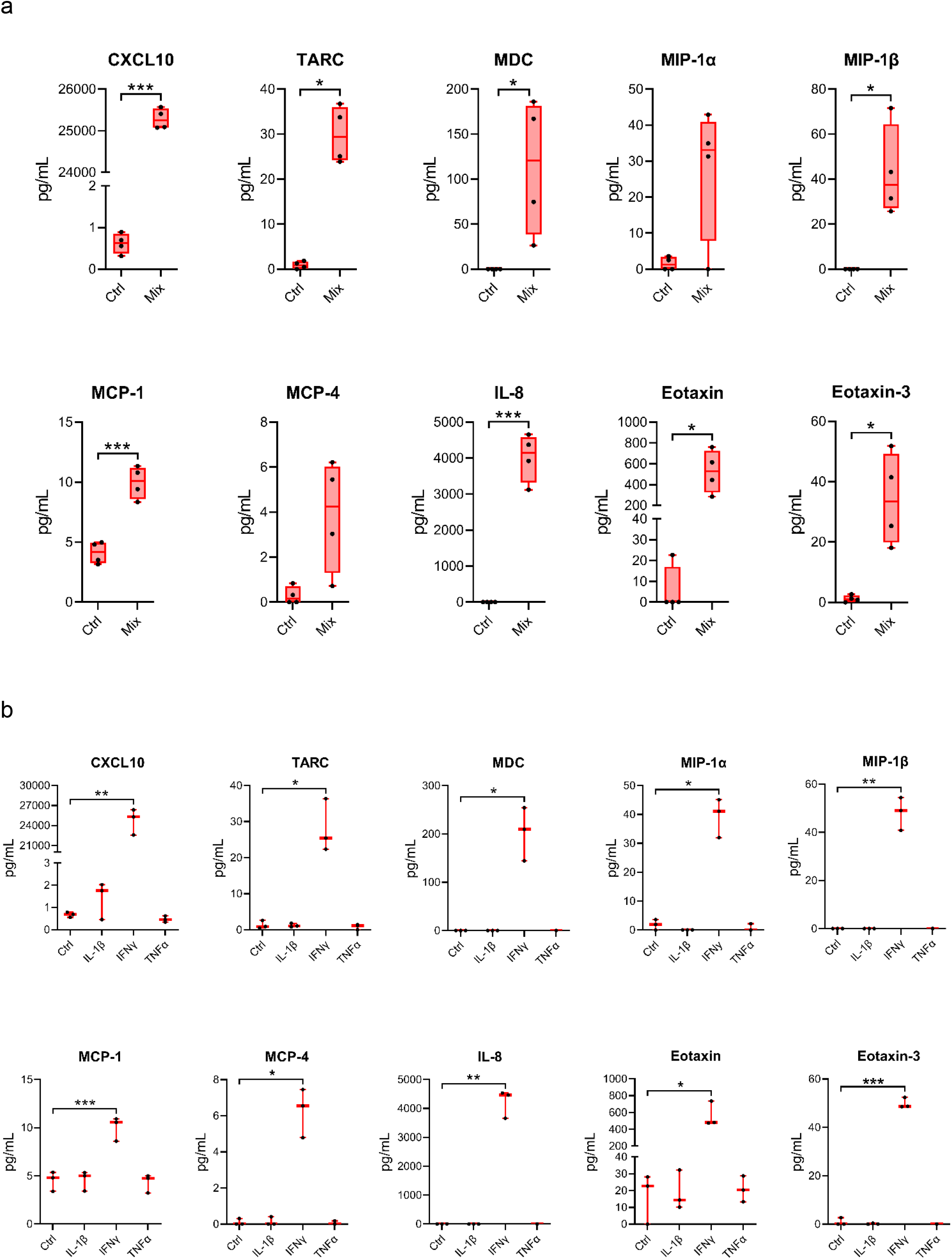
Cytokine-induced chemokine secretion from EndoC-βH5 cells. Accumulated chemokines in the culture medium from EndoC-βH5 cells after 48 h of cytokine treatment (50 U/mL IL-1β, 1000 U/mL IFNγ and 1000 U/mL TNFα) in combination (Mix) (a) or individual cytokines (b), as compared to untreated cells. Data are mean pg/mL with median and 5/95 percentiles (n=3-4), *p<0.05, **p<0.01, ***p<0.001.

### Insulin secretion from cytokine-exposed EndoC-βH5 cells

To investigate the impact of cytokines on the insulin secretory capacity of EndoC-βH5 cells, GSIS experiments were performed following exposure to IL-1β, IFNγ and TNFα for 48 hours. In untreated control cells, there was a significant increase in GSIS corresponding to a fold change of >6 (Fig. 6a). This induction was diminished by ~30% by cytokine exposure (Fig. 6a). This effect correlated with tendencies towards reduced cellular insulin content (Fig. 6b), decreased expression of *INS* (Fig. 6c), and decreased *PDX1* expression (Fig. 6d). These results demonstrate that cytokines cause functional impairment of EndoC-βH5 cells.

**Figure 6:**
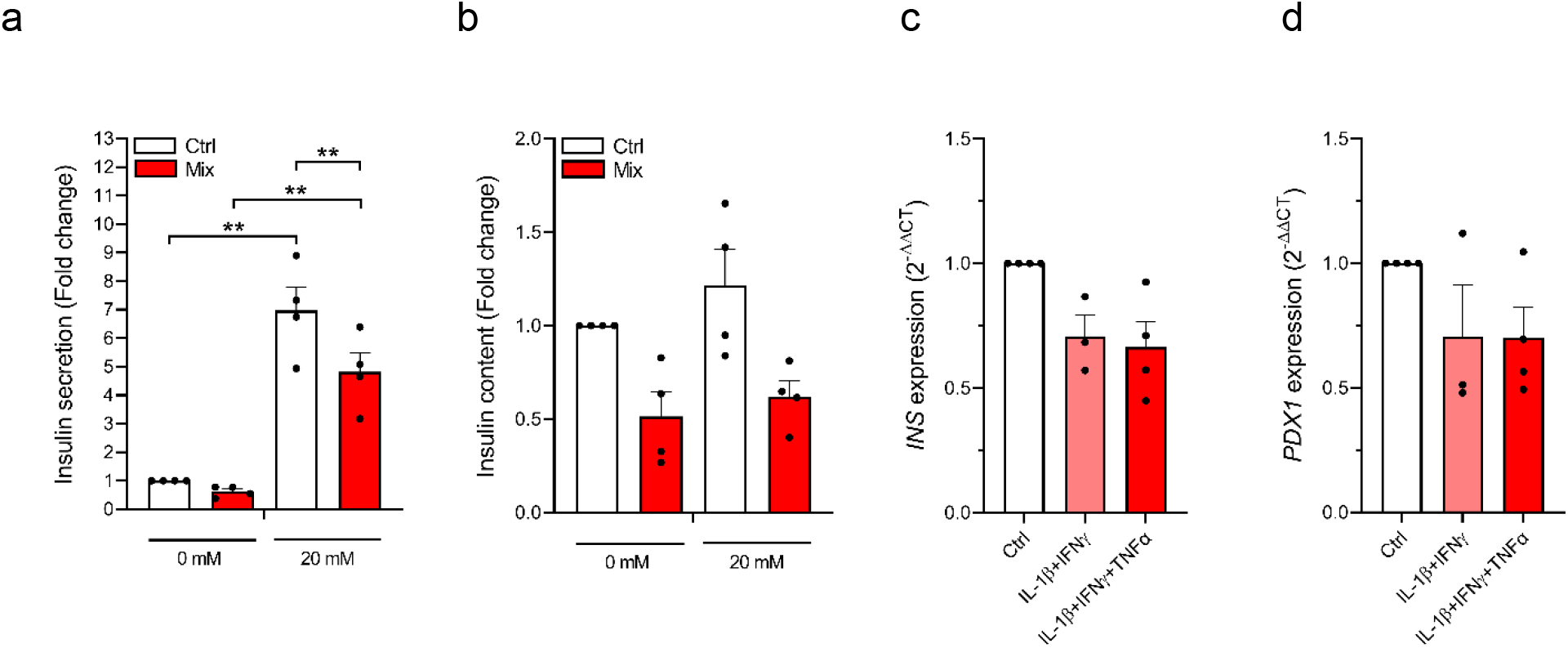
Cytokine-induced functional impairment of EndoC-βH5 cells. Glucose-stimulated insulin secretion (a) and insulin content (b) after 48 h of cytokine treatment (50 U/mL IL-1β, 1000 U/mL IFNγ and 1000 U/mL TNFα) (Mix) in response to 0 mM or 20 mM glucose stimulation. Data was normalised to DNA content. Gene expression of *INS* (c) and *PDX1* (d) in EndoC-βH5 cells after 48 h of cytokine treatment. *GAPDH* was used as housekeeping gene. Data are means ± SEM (n=3-4), **p<0.01.

### Mitochondrial respiration in cytokine-exposed EndoC-βH5 cells

To investigate the effects of cytokines on mitochondrial function of EndoC-βH5 cells, we treated cells with individual or combinations of cytokines for 48 hours followed by measurement of oxygen consumption rate (OCR) using Seahorse extracellular flux technology (Fig. 7a). Basal respiration decreased after cytokine exposure in all conditions except IL-1β alone (Fig. 7b). Glucose oxidation, proton leak and ATP production were decreased in response to treatment with a combination of the cytokines. Glucose response was enhanced in cells exposed to the combined cytokines, as compared to control (Fig. 7c). The results suggest that cytokines, and particularly IFNγ, perturb mitochondrial respiration in EndoC-βH5 cells.

**Figure 7:**
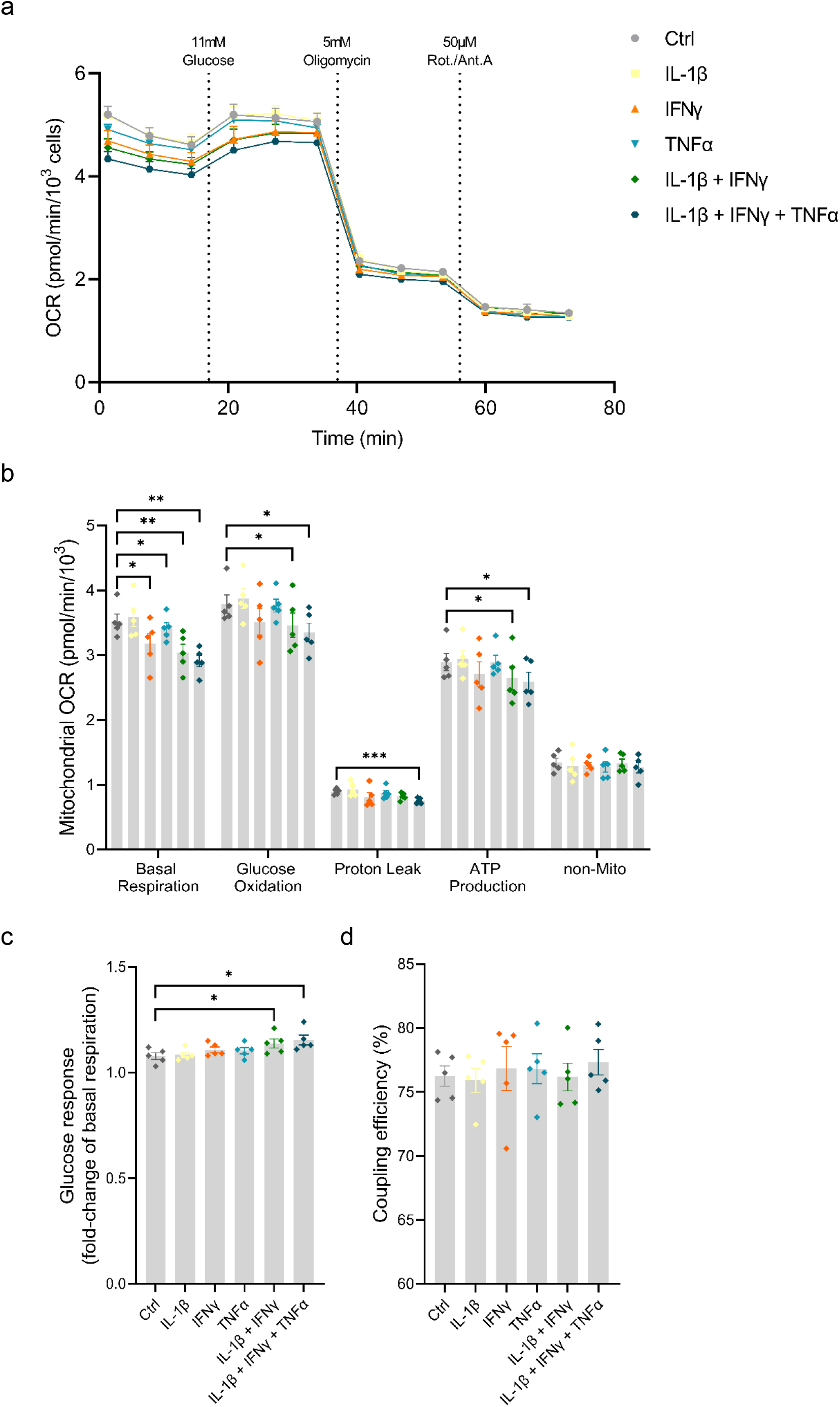
Cytokines impair mitochondrial respiration in EndoC-βH5 cells. Mitochondrial respiration in untreated cells and cells exposed to cytokines (50 U/mL IL-1β, 1000 U/mL IFNγ and l000 U/mL TNFα) alone or in combinations, as indicated, for 48 h. (a) OCR traces normalised to cell number per well. (b) Analysis of mitochondrial parameters from OCR traces were corrected for non-mitochondrial respiration. (c) Glucose response of respiration as fold-change of basal mitochondrial respiration. (d) Coupling efficiency calculated as the oligomycin-sensitive fraction of glucose-stimulated mitochondrial respiration. Results are means ± SEM (n=5), *p<0.05, **p<0.01, ***p<0.001.

### Cytokine effects on the transcriptome of EndoC-βH5 cells

We next studied the impact of cytokines on the global gene expression profile of EndoC-βH5 cells. Cells were treated with or without IL-1β, IFNγ and TNFα for 48 hours followed by stranded RNA sequencing. On average, library sizes of ~40 million reads per sample were obtained (Fig. 8a). Multidimensional scaling (MDS) analysis showed a high separation between control and cytokine-treated samples (Fig. 8b). A total of 16,525 genes were detected of which almost two-thirds (6,000 genes) were differentially expressed in response to cytokines (Fig. 8c; cut off of abs(log2FC) >0.5 and FDR adjusted p-value <0.05). The top 10 up- and downregulated coding (mRNA) and lncRNA genes are shown in Table 1. The data obtained demonstrate that cytokine treatment causes profound changes to the transcriptome of EndoC-βH5 cells.

**Figure 8:**
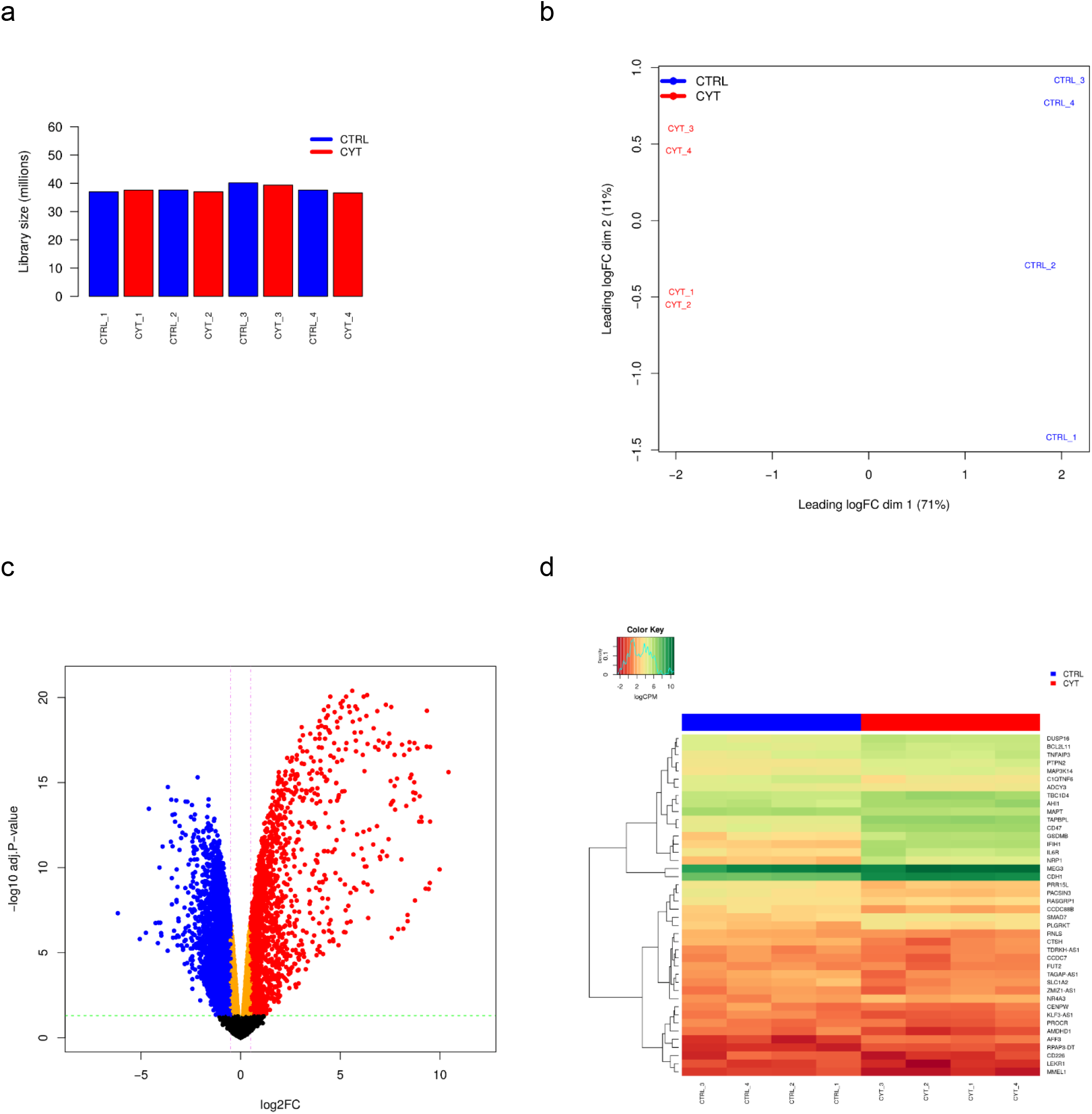
Cytokines modulate a large part of the gene expression profile of EndoC-βH5 cells. RNA sequencing of untreated control cells or cells exposed to cytokines (50 U/mL IL-1β, 1000 U/mL IFNγ and 1000 U/mL TNFα) for 48 h. (a) Library size of each of the RNA samples sequenced (CTRL_1-4, 4 replicates of untreated control cells; CYT_1-4, 4 replicates of cytokine-exposed cells). (b) MDS plot and (c) Volcano plot showing differentially expressed genes after exposure to cytokines compared to untreated cells. The horizontal line represents the adjusted p-value and the vertical line represent fold change cut-offs. The red and blue dots represent the upregulated and downregulated genes, respectively. (d) Heat map and hierarchical clustering of the 42 differentially expressed candidate genes mapping to type 1 diabetes loci.

**Table 1.** Ten most up- and downregulated mRNAs and lncRNAs ranked based on log-fold changes (logFC).

| mRNAs | Gene ID | Gene name | Chr. | logFC | p-value | FDR |
| --- | --- | --- | --- | --- | --- | --- |
| 10 most upregulated | ENSG00000177409 | *SAMD9L* | 7 | 10.4276 | 1.41<sup>-18</sup> | 2.42<sup>-16</sup> |
|  | ENSG00000136514 | *RTP4* | 3 | 9.980989 | 4.46<sup>-12</sup> | 1.27<sup>-10</sup> |
|  | ENSG00000169245 | *CXCL10* | 4 | 9.504668 | 2.8<sup>-15</sup> | 1.96<sup>-13</sup> |
|  | ENSG00000156219 | *ART3* | 4 | 9.433 | 9.74<sup>-11</sup> | 1.81<sup>-09</sup> |
|  | ENSG00000169248 | *CXCL11* | 4 | 9.343853 | 5.83<sup>-23</sup> | 6.03<sup>-20</sup> |
|  | ENSG00000090339 | *ICAM1* | 19 | 9.336642 | 2.68<sup>-20</sup> | 7.64<sup>-18</sup> |
|  | ENSG00000130303 | *BST2* | 19 | 9.077847 | 2.82<sup>-15</sup> | 1.97<sup>-13</sup> |
|  | ENSG00000162645 | *GBP2* | 1 | 8.983403 | 6.28<sup>-17</sup> | 6.33<sup>-15</sup> |
|  | ENSG00000117228 | *GBP1* | 1 | 8.957278 | 2.84<sup>-15</sup> | 1.97<sup>-13</sup> |
|  | ENSG00000173110 | *HSPA6* | 1 | 8.872981 | 3.78<sup>-20</sup> | 9.76<sup>-18</sup> |
| 10 most downregulated | ENSG00000267909 | *CCDC177* | 14 | -4.74879 | 9.57<sup>-08</sup> | 6.77<sup>-07</sup> |
|  | ENSG00000164363 | *SLC6A18* | 5 | -4.26357 | 2.06<sup>-08</sup> | 1.79<sup>-07</sup> |
|  | ENSG00000100079 | *LGALS2* | 22 | -4.14731 | 4.77<sup>-07</sup> | 2.72<sup>-06</sup> |
|  | ENSG00000086967 | *MYBPC2* | 19 | -4.12385 | 1.95<sup>-08</sup> | 1.71<sup>-07</sup> |
|  | ENSG00000126861 | *OMG* | 17 | -4.02992 | 1.97<sup>-08</sup> | 1.72<sup>-07</sup> |
|  | ENSG00000205022 | *PABPN1L* | 16 | -3.98799 | 1<sup>-07</sup> | 7.04<sup>-07</sup> |
|  | ENSG00000187783 | *TMEM72* | 10 | -3.97659 | 1.46<sup>-07</sup> | 9.83<sup>-07</sup> |
|  | ENSG00000154134 | *ROBO3* | 11 | -3.91304 | 1.27<sup>-13</sup> | 5.7<sup>-12</sup> |
|  | ENSG00000135312 | *HTR1B* | 6 | -3.65834 | 1.58<sup>-06</sup> | 7.71<sup>-06</sup> |
|  | ENSG00000183379 | *SYNDIG1L* | 14 | -3.64772 | 1.49<sup>-17</sup> | 1.82<sup>-15</sup> |
| lncRNAs | Gene ID | Gene name | Chr. | logFC | p-value | FDR |
| 10 most upregulated | ENSG00000282851 | *BISPR* | 19 | 9.503658 | 2.84<sup>-20</sup> | 7.97<sup>-18</sup> |
|  | ENSG00000224666 | *ETV7-AS1* | 6 | 9.490858 | 3.69<sup>-11</sup> | 7.9<sup>-10</sup> |
|  | ENSG00000289582 | – | 1 | 9.28052 | 9.19<sup>-11</sup> | 1.72<sup>-09</sup> |
|  | ENSG00000206337 | *HCP5* | 6 | 7.748361 | 4.79<sup>-16</sup> | 3.9<sup>-14</sup> |
|  | ENSG00000256262 | *USP30-AS1* | 12 | 7.521434 | 5.92<sup>-18</sup> | 8.09<sup>-16</sup> |
|  | ENSG00000272941 | – | 7 | 6.932371 | 9.69<sup>-21</sup> | 3.37<sup>-18</sup> |
|  | ENSG00000273272 | – | 22 | 6.399633 | 9.34<sup>-09</sup> | 9.14<sup>-08</sup> |
|  | ENSG00000261618 | *LINC02605* | 8 | 6.15968 | 1.34<sup>-14</sup> | 7.69<sup>-13</sup> |
|  | ENSG00000286647 | – | 5 | 6.155537 | 9.57<sup>-13</sup> | 3.32<sup>-11</sup> |
|  | ENSG00000261644 | *CYLD-AS1* | 16 | 6.064333 | 1.76<sup>-13</sup> | 7.47<sup>-12</sup> |
| 10 most downregulated | ENSG00000261240 | – | 16 | -6.15617 | 4.45<sup>-09</sup> | 4.77<sup>-08</sup> |
|  | ENSG00000260573 | – | 16 | -5.05595 | 2.53<sup>-07</sup> | 1.57<sup>-06</sup> |
|  | ENSG00000249201 | *TERLR1* | 5 | -4.60144 | 4.18<sup>-16</sup> | 3.44<sup>-14</sup> |
|  | ENSG00000246985 | *SOCS2-AS1* | 12 | -4.06762 | 3.37<sup>-12</sup> | 9.86<sup>-11</sup> |
|  | ENSG00000227487 | *NCAM1-AS1* | 11 | -3.3381 | 6.98<sup>-08</sup> | 5.14<sup>-07</sup> |
|  | ENSG00000225649 | – | 2 | -3.28783 | 2.89<sup>-15</sup> | 1.99<sup>-13</sup> |
|  | ENSG00000266036 | *SLC9A3R1-AS1* | 17 | -3.21135 | 5.51<sup>-10</sup> | 7.97<sup>-09</sup> |
|  | ENSG00000224186 | *PITX1-AS1* | 5 | -3.16412 | 1.44<sup>-07</sup> | 9.71<sup>-07</sup> |
|  | ENSG00000272549 | *LINC02538* | 6 | -2.96695 | 6.8<sup>-07</sup> | 3.7<sup>-06</sup> |
|  | ENSG00000130600 | *H19* | 11 | -2.91744 | 1.58<sup>-14</sup> | 8.87<sup>-13</sup> |

We extracted gene expression data of pinpointed candidate genes from 152 type 1 diabetes-associated loci and identified 42 candidate genes that were differentially expressed upon cytokine exposure. Twenty genes were significantly upregulated and 22 genes significantly downregulated (Fig. 8d).

### Pathway analysis of differentially expressed genes

Using IPA, we were able to predict the top relevant disease-related and molecular interactions of the differentially expressed genes after exposure to cytokines. The Core analysis revealed that the top affected canonical pathway was the Senescence Pathway, and the top affected molecular and cellular function was Cell Death and Survival (Table 2). Most significant upstream regulators, diseases, and mechanisms of the differentially expressed genes are represented in a graphical summary network (Fig. 9). Activated pathways in this network included Diabetes mellitus, Senescence Pathway, Endocrine pancreatic dysfunction, Interferon Signalling and Organismal death.

**Table 2.** Top canonical pathways and top molecular and cellular functions predicted for the differentially expressed genes from the RNA sequencing data using Ingenuity Pathway Analysis (IPA). The table shows the p-value or p-value range, as well as the overlapping targets identified for each pathway/function, as predicted by the Core analysis in the IPA software.

| Top canonical pathways | p-value | Overlap |
| --- | --- | --- |
| Senescence Pathway | $1.28 \times 10^{-9}$ | 34.4% (103/299) |
| Kinetochore Metaphase Signalling Pathway | $1.27 \times 10^{-8}$ | 43.2% (48/111) |
| Molecular Mechanisms of Cancer | $9.04 \times 10^{-8}$ | 30.0% (135/450) |
| Axonal Guidance Signalling | $1.09 \times 10^{-7}$ | 29.3% (149/509) |
| Protein Kinase A Signalling | $2.12 \times 10^{-7}$ | 30.2% (124/411) |
| Molecular and Cellular Functions | p-value range | Molecules |
| Cell Death and Survival | $1.17 \times 10^{-10} - 3.16 \times 10^{-33}$ | 1566 |
| Cellular Assembly and Organization | $1.12 \times 10^{-12} - 4.21 \times 10^{-29}$ | 827 |
| Cellular Function and Maintenance | $8.37 \times 10^{-16} - 4.21 \times 10^{-29}$ | 1317 |
| Cellular Movement | $8.07 \times 10^{-11} - 1.20 \times 10^{-26}$ | 1157 |
| Cellular Development | $3.83 \times 10^{-11} - 3.05 \times 10^{-26}$ | 1640 |

**Figure 9:**
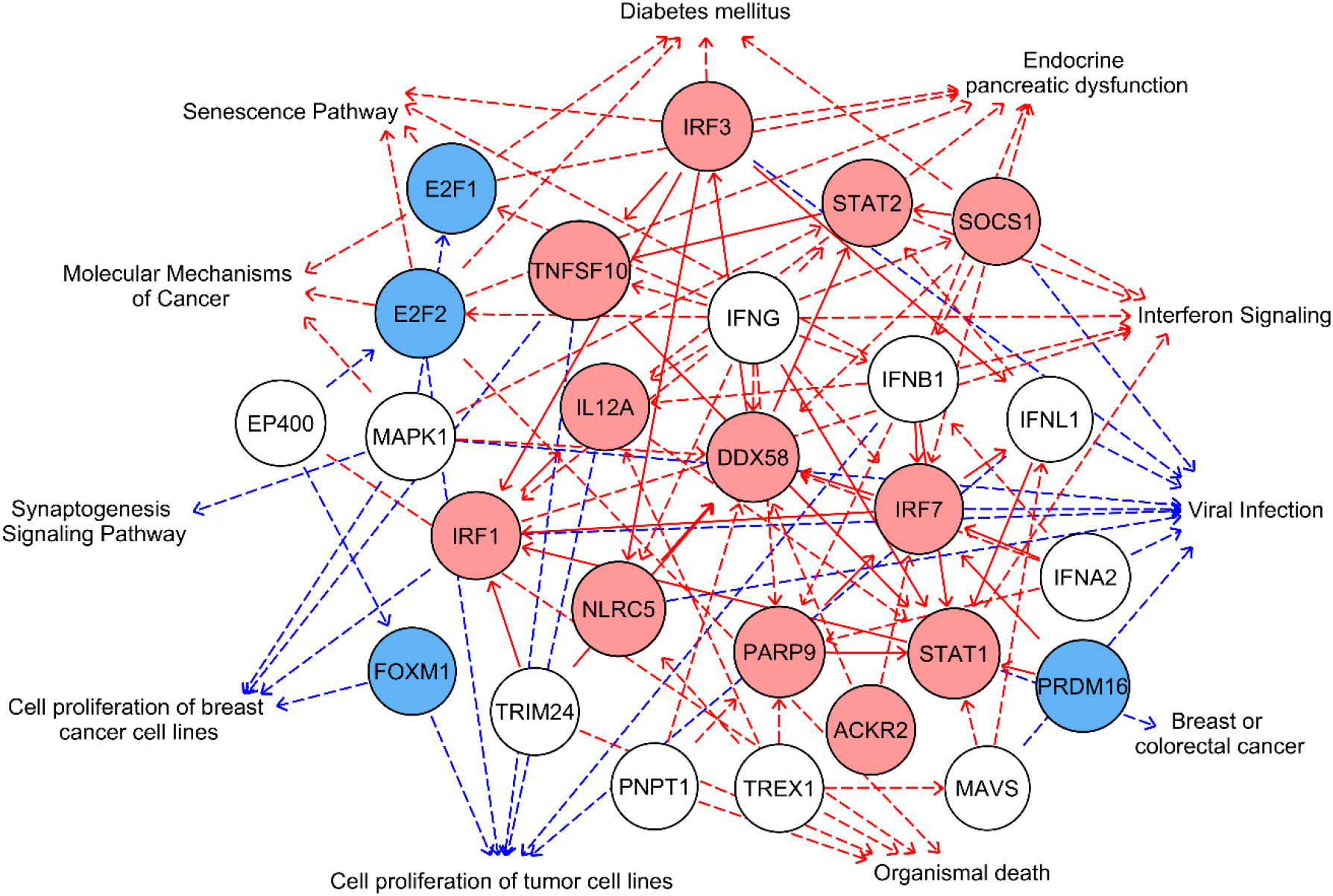
Summary network of the differentially expressed genes in cytokine-treated EndoC-βH5 cells. Graphical summary network representing the most significant upstream regulators, diseases, functions, and pathways, and their relations, as predicted by the IPA Core analysis. Red edges: predicted activated pathways, blue edges: predicted inhibited pathways. Solid lines represent direct relationships and dashed lines represent indirect relationships. Red nodes: upregulated, blue nodes: downregulated, white nodes: non-affected, based on differential expression from dataset.

### Expression of key lncRNAs in EndoC-βH5 cells

As lncRNAs are emerging as important regulators of cellular functions and disease, we wished to validate the expression of selected lncRNAs by qPCR. EndoC-βH5 cells exposed to cytokines for 48 hours showed significantly decreased expression of the lncRNAs *TUG1* and *TUNAR* (Fig. 10a and b). The lncRNAs *MEG3* and *GAS5* were not significantly affected by the cytokine exposure (Fig. 10c and d). *LINC-25* was not expressed (data not shown).

**Figure 10:**
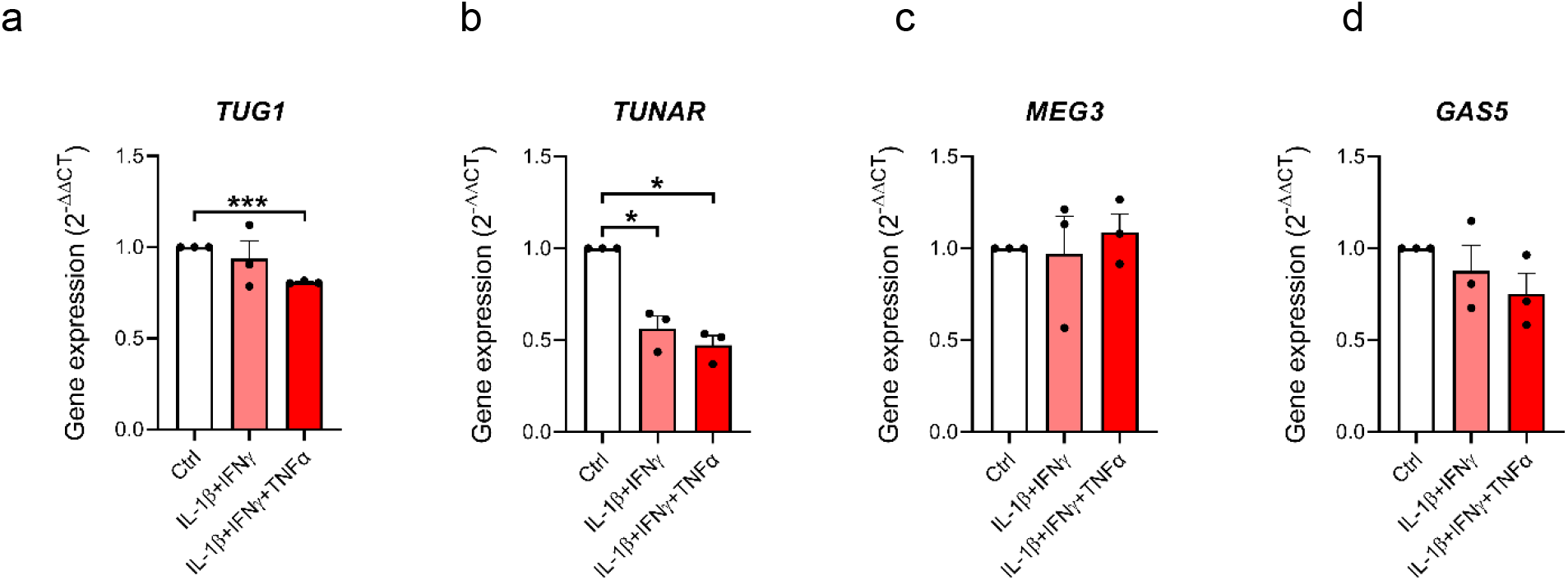
LncRNA expression in EndoC-βH5 cells. Expression of 4 lncRNAs (a) *TUG1,* (b) *TUNAR,* (c) *MEG3* and (d) *GAS5* after exposure to cytokines (50 U/mL IL-1β, l000 U/mL IFNγ and 1000 U/mL TNFα) for 48 h. *GAPDH* was used as housekeeping gene. Data are means ± SEM (n=3), *p<0.05, ***p<0.001.

## Discussion

In the present study, we characterised the sensitivity profile and cellular responses of human EndoC-βH5 cells to diabetogenic cytokines which is a widely used in vitro model of beta-cell destruction in type 1 diabetes.

The observed time- and dose-dependent increase in caspase 3/7 activity and cytotoxicity in EndoC-βH5 cells in response to cytokines is in agreement with studies on EndoC-βH1 cells [5] and human pancreatic islets [24]. Interestingly, our study identified IFNγ as the sole pro-apoptotic cytokine. As shown by immunoblotting, the level of the downstream mediator of IFNγ, P-STAT1, was significantly elevated by IFNγ, whereas P-JNK and IκBα downstream of IL-1β and TNFα receptor signalling, were unaffected by cytokine treatment. Thus, IFNγ seems to be the main driver of signalling and apoptosis in EndoC-βH5 cells in the time window examined. This contrasts with what has been reported for EndoC-βH1 cells, which shows that IL-1β alone and to a lesser extent TNFα alone increases apoptosis measured by caspase 3/7 activity [5]. The importance of IFN as a potential driving diabetogenic cytokine was recently highlighted in a study by Apaolaza and colleagues, who investigated the IFN signature in islets from autoantibody-positive human donors with type 1 diabetes [26]. The study reported that IFN response markers colocalised with insulin-containing cells in islets with insulitis from donors with type 1 diabetes and autoantibodies, as compared to healthy control islets. Several studies in mice have also stressed IFNγ as a key player in insulitis, being essential for beta-cell targeting and the development of diabetes [27–29]. Further studies aimed at defining the pathophysiological role of IFNγ in type 1 diabetes are warranted and may identify rational points of intervention to halt beta-cell killing.

By TUNEL assay, we observed a basal cell death rate for EndoC-βH5 cells of around 20%, which may foremost represent the non-viable cell fraction observed following thawing of the cells. This rate is higher as compared to what is typically observed for other beta-cell models, which have basal death rates of 5-10% [15, 30, 31]. This may explain the rather modest ~2-fold cytokine effect on cell death observed in EndoC-βH5 cells. The higher basal death might reflect a more fragile phenotype of the EndoC-βH5 cells compared to other beta-cell lines, which are immortalized and generally exhibit a less differentiated phenotype, including the EndoC-βH1 cell line [30]. For human islets, basal cell death varies significantly among studies, between 10-20% [31–33]. This may reflect that EndoC-βH5 cells resemble primary human beta cells more closely than other beta-cell lines.

A key characteristic of cytokine-exposed beta cells is the upregulation of immune-modulatory proteins and antigen presentation [21]. The MHC class I molecules (encoded by *HLA)* play a pivotal role in the recognition of beta cells by autoreactive T cells. We showed that MHC-I was upregulated by cytokines at both the protein and mRNA level. Furthermore, several other beta-cell-produced factors have immune-modulatory effects including chemoattraction and inflammation [14]. Accordingly, we show that the EndoC-βH5 cells secreted several chemokines in response to cytokines. This contributes to their diabetogenic phenotype following cytokine exposure and highlight the use of the EndoC-βH5 cells as a valid model to study the beta cell-immune cell crosstalk in terms of chemokine production. From single cytokine exposure experiments, we found that chemokine secretion was exclusively driven by IFNγ, which verifies our previous findings, that the diabetogenic effects arise from the IFNγ exposure.

One of the main reasons for choosing EndoC-βH5 cells over other beta-cell models is their reported higher insulin content and superior insulin secretory capacity [1, 5, 6, 34]. We found that upon stimulation with high glucose, the EndoC-βH5 cells responded with a more than 6-fold induction in insulin release compared to low glucose, which is in agreement with recently published studies [10, 11]. The insulin secretory capacity was diminished by cytokine exposure, in accordance with findings in human and rodent beta-cell lines and isolated pancreatic islets [5, 35–38]. The cytokine-mediated blunted insulin secretion was observed after 48 hours of cytokine exposure i.e., at a time point before increased cell death was observed, underlining that cytokine-induced functional impairment precedes the induction of cell death.

Our data on mitochondrial function showed reduced respiration following treatment with cytokines. Surprisingly, on several mitochondrial parameters including basal respiration, glucose oxidation, proton leak, and ATP production, combinations of cytokines had the most pronounced inhibitory effects. This observation suggests some degree of synergism between the cytokines with regard to mitochondrial impairment. The underlying mechanisms of cytokine-mediated impaired mitochondrial function are currently unclear but would be relevant to address in future studies. We observed that the glucose response was slightly but significantly increased in EndoC-βH5 cells after exposure to cytokine combinations. This may reflect an increased level of oxidative stress, which has been shown to be involved in beta-cell dysfunction [39].

Around 50% of the genetic risk of type 1 diabetes resides within the *HLA* region [40]. The remaining risk loci harbours both protein-coding and non-coding genomic regions, emphasising the implication of both in type 1 diabetes pathogenesis [19, 40]. By RNA-seq analysis, we found that 42 pinpointed candidate risk genes from type 1 diabetes-associated loci were differentially expressed after cytokine exposure. These included both mRNAs and lncRNAs. Overall, we detected more than 16,000 genes expressed in EndoC-βH5 cells of which more than a third (6,000 genes: 1,934 up; 4,066 down) were modified by cytokines. This differential expression is seemingly larger than what was previously reported in similar studies of human islets and Endo-βH1 cells [41, 42], and greatly reflect the large number of downregulated genes that we report here. These observations may represent phenotypic variations but may also result from overall experimental differences. By pathway analysis, we were able to predict relevant molecular interactions, including functions in Cell Death and Survival, and within Diabetes. Furthermore, we validated several mRNAs and lncRNAs that are known to be implicated in beta-cell apoptosis and dysfunction [21, 22].

Due to the lack of evidence for functional and signalling effects of IL-1β and TNFα, the presence of their receptors in the cells could be questioned. Retrieval of the CPM values of the IL-1 and TNFα receptors from the RNA-seq data revealed that these were lowly expressed (Supplementary Table S1) – a finding that was also confirmed by qPCR (data not shown). In contrast, the IFNγ receptor was expressed at a much higher level. This is a plausible explanation as to why IL-1β and TNFα failed to show individual effects or synergistic effects when combined with IFNγ. This is an important aspect and possible drawback of the EndoC-βH5 cells that should be considered when using this model for studying cytokine effects. It also highlights the importance of performing thorough characterisation of model systems in general.

Thus far, only few studies have been published on EndoC-βH5 cells [10, 11], but the cells hold promise as a superior model of native human beta cells over other human beta-cell models. The validity of the human hybrid beta-cell line 1.1B4 [43] must be questioned, as a study reported rodent cell contamination in the cell stocks and established deprivation of insulin and glucose responsiveness after isolation of the human cell population [34]. The EndoC-βH1 cell line is the most widely used human beta-cell line and has been used by many research laboratories for the past 10 years [4–6, 44]. However, EndoC-βH1 cells suffer from functional limitations compared to native beta cells [5–9]. Moreover, during the original expansion of the EndoC-βH1 cells in mice, the cells were stably infected with a xenotropic murine virus [7]. It is still uncertain to which extent the infection with this virus affects the cells’ phenotype and cautions should be taken when extrapolating results from this cell line.

Although of relevance, we chose not to perform a direct head-to-head comparison of the EndoC-βH5 cells with other cell models. The present work is therefore limited with regards to determining the exact discrepancies between EndoC-βH5 cells and e.g., human islets and EndoC-βH1 cells. However, our study offers detailed insight into the EndoC-βH5 cells’ cytokine sensitivity profile and cellular responses including functional and transcriptomic effects. Besides being advantageous by having non-cancerous properties, i.e., a non-proliferative phenotype, and absence of the xenotropic murine virus, EndoC-βH5 cells possess key beta-cell properties including a high insulin secretory capacity. Our study adds to this by demonstrating that IFNγ, a key type 1 diabetogenic cytokine, induces several relevant cellular responses including upregulation of MHC-I, STAT1 activation, and chemokine secretion. To our knowledge, this is the first study describing the EndoC-βH5 cells’ sensitivity profile to cytokines. The cells’ usage for cytokine studies may be limited by their apparent exclusive responsiveness to IFNγ, possibly due to a very low expression level of the IL-1 and TNFα receptors. It will be important to further investigate the EndoC-βH5 cells’ response and behaviour to other diabetogenic stimuli and to directly compare them with primary human beta cells.

## Abbreviations

GSIS: glucose-stimulated insulin secretion
IL-1β: interleukin-1β
IFNγ: interferon γ
IκBα: nuclear factor of kappa light polypeptide gene enhancer in B-cells inhibitor α
IPA: Ingenuity Pathway Analysis
lncRNA: long non-coding RNA
MDS: multidimensional scaling
MHC: major histocompatibility complex
OCR: oxygen consumption rate
P-JNK: phosphor-c-Jun N-terminal kinase
P-STAT1: phospho-signal transducer and activator of transcription 1
RNA-seq: RNA sequencing
TNFα: tumor necrosis factor-α

## Data availability

The RNA-seq data presented in this study can be found online: GEO assession number GSE218735.

## Authors’ relationships and activities

The authors declare that there are no relationships or activities that could have influenced the work performed in this study.

## Contribution statement

C.F. and J.S. conceptualized the study. C.F., R.H.G., F.P., S.K. and J.S. designed the study. C.F., R.H.G., C.A.S.S., J.M.L.M., T.F. and S.K. contributed to the acquisition of data. C.F. made the first draft of the manuscript which was commented by J.S. All authors contributed with critical scientific input, interpretation of data and manuscript writing. The final version of the manuscript was approved by all authors.

