## Supplementary Table S1 for "Characterisation of the functional and transcriptomic effects of pro-inflammatory cytokines on human EndoC-βH5 beta cells"

Supplementary materials

**Supplementary Table S1** CPM values of cytokine receptors retrieved from RNA sequencing of untreated control cells or cells exposed to cytokines (50 U/mL IL-1β, 1000 U/mL IFNγ and 1000 U/mL TNFα) for 48 h (CTRL_1-4, 4 replicates of untreated control cells; CYT_1-4, 4 replicates of cytokine-exposed cells).

| **Gene ID** | **Gene name** | **Chr.** | **1-CTRL** | **2-CTRL** | **3-CTRL** | **4-CTRL** | **1-CYT** | **2-CYT** | **3-CYT** | **4-CYT** |
|---|---|---|---|---|---|---|---|---|---|---|
| ENSG00000027697 | *IFNGR1* | 6 | 26.54 | 28.34 | 31.71 | 26.24 | 33.58 | 33.00 | 33.08 | 34.56 |
| ENSG00000159128 | *IFNGR2* | 21 | 50.14 | 56.05 | 76.48 | 63.39 | 49.50 | 49.98 | 75.35 | 61.10 |
| ENSG00000159110 | *IL1R1* | 2 | 0.86 | 0.61 | 0.48 | 0.57 | 0.96 | 1.43 | 0.48 | 0.42 |
| ENSG00000185436 | *TNFRSF1B* | 1 | 0.00 | 0.00 | 0.00 | 0.00 | 0.70 | 0.23 | 0.62 | 0.42 |
